# Plasma Extracellular Vesicle and miRNA Profiles in Pregnancies Complicated by Fetal Congenital Heart Defects

**DOI:** 10.64898/2026.05.23.727381

**Authors:** Alyssa Tipler, Marcella Rodriquez, Mayu Morita, Stephanie Y. Tseng, James Cnota, Terry K Morgan, Helen N. Jones

## Abstract

Congenital heart defects (CHDs) are the most common form of fetal malformation however, our understanding of trophoblast health and communication throughout gestation in CHD pregnancies remains limited. The purpose of this study was to assess extracellular vesicles (EVs) and microRNA (miRNA) present in maternal and umbilical cord plasma from a spectrum of CHD subtypes during gestation and at time of delivery. We hypothesized that circulating placenta-derived EVs and miRNA will differ in CHD when compared to controls. Maternal plasma samples were collected between 16-24 weeks of gestation and at the time of delivery. Umbilical cord plasma was obtained following delivery. EVs were isolated from plasma samples using nanoscale flow cytometry, and total EV counts as well as counts by cellular origin were determined. MicroRNA was extracted from maternal plasma and levels quantified using qPCR. Maternal plasma from pregnancies complicated by fetal CHD exhibited higher total EV counts at delivery compared to control. Platelets derived extracellular vesicles (pdEVs) were significantly higher both in maternal and cord blood plasma at the time of delivery in CHD pregnancies compared to gestationally age-matched control pregnancies. Circulating miR22 and miR421 levels were reduced, while miR29c levels were increased in maternal plasma from CHD pregnancies between 16-24 weeks but no differences seen at time of delivery. Pregnancies complicated by CHD are associated with an altered in utero environment by changes in extracellular vesicles and miRNA profile in maternal serum. Circulating EVs and miRNA profiles may therefore serve as minimally invasive indicators of placental and maternal vascular dysfunction in CHD.

## INTRODUCTION

Congenital heart defects (CHDs) are the most common form of birth anomalies, affecting nearly 1% of all live births annually^1^. The structural abnormalities of the heart occur during fetal development and can range in severity from minor defects, requiring no intervention, to complex malformations that require surgery. Recent evidence highlights the critical role of the placenta in fetal cardiovascular development, growth and remodeling as it regulates nutrient and oxygen availability ^2^ however, its role is poorly understood. This study characterizes circulating cell-type specific EVs and miRNAs in maternal blood at the prenatal visit and umbilical cord blood at delivery across various CHD subtypes to identify cell to cell communication pathways. We hypothesize that circulating placenta-derived EVs and miRNA will be different in CHD compared to controls offering a mechanism to assess placental health and develop future interventions.

Extracellular vesicles are small, membrane-bound vesicles released by cells into the extracellular environment that play a critical role in intercellular communication by transferring cargoes of bioactive molecules including proteins, lipids, and RNA^3^. In pregnancy, EVs are released from the extravillous trophoblast and syncytiotrophoblast in addition to maternal cells (endothelial cells, platelets, immune cells) into the maternal circulation and can be detected as early as 6 weeks of gestation^4^. As gestation continues, EV abundance increases, allowing for the assessment of the longitudinal health of the placenta in maternal blood samples^4,5^. EV membranes contain specific markers that can be used to determine their cell of origin, potentially allowing for cell-specific assessment^6^. Studies demonstrated that placenta-derived EVs can be distinguished by plasma membrane markers such as Placental Alkaline Phosphatase (PLAP), which is enriched in the microvillous membrane of the syncytiotrophoblast^7^. In pregnancies complicated by gestational diabetes mellitus (GDM)^8^ and preeclampsia^9^ there is an increase in total and placental-derived EVs in maternal plasma at the time of delivery. Assessing the cargo within these EVs, such as miRNA helps us to better understand cell to cell communication between placenta and maternal cells. Although EVs membrane markers can be used to identify the cellular origin, current technical limitations restrict the ability to isolate purified sorted EV populations for downstream miRNA cargo analysis. Therefore, the miRNA content assessed in this study represent total circulating EV-associated miRNA content rather than cargo derived from a specific EV cell population, thus limiting the ability to determine their precise cellular source.

MicroRNAs (miRNAs) are 20-25 nucleotide non-coding RNA molecules that play a crucial role in post-transcriptional regulation of gene expression^10^. These miRNAs often regulate gene expression by binding to mRNA and either promoting their degradation or inhibiting translation. In the placenta, miRNAs are highly abundant and help regulate is critical processes including trophoblast proliferation, differentiation invasion and vascular remodeling^11^. In the maternal blood, placental miRNAs circulate within EVs where they modulate maternal physiology, immune responses and endothelial function to support pregnancy. In this study, we focus on evaluating the levels of miR-22, miR-29c, and miR-421 in maternal blood samples. Studies have demonstrated increased levels of these miRNAs in second trimester maternal blood in pregnancies affected by specific CHDs, including ventricular septal defects (VSD), atrial septal defects (ASD) and tetralogy of Fallot (TOF) ^12^. We evaluated the levels of these miRNAs in various CHD subtypes at 20±4 weeks and at the time of delivery.

Based on the intertwined connections between placental dysfunction and fetal cardiovascular defects, we aim to analyze maternal blood samples for the presence of EVs and miRNAs as these parameters may provide insight into the health of the placenta. We hypothesize that circulating placenta-derived EVs and miRNAs will be different in CHD samples compared to controls, offering a mechanism to assess placental health and develop future interventions. This could potentially advance our understanding of cell to cell communication and provide a non-invasive method for monitoring pregnancies complicated by CHD.

## MATERIALS AND METHODS

### Patient Enrollment and Sample Collection

This study utilized a cohort assembled across two centers: University of Florida Health Women’s Health Center – Medical Plaza (IRB #202101799) and Cincinnati Children’s Hospital Medical Center (IRB #2018-8095). Written informed consent was obtained from all participants in accordance with protocols approved by the Institutional Review Boards at each institution. The study included pregnant patients aged 18–45 years carrying fetuses at risk for congenital heart disease (CHD), as identified through fetal sonographic screening, genetic screening results, family history, or maternal risk factors. Exclusion criteria included pregnancies resulting from assisted reproductive technologies, multiple gestations, mothers with known pregestational diabetes, and mothers with corrected CHD or other known structural heart disease. For samples collected at Cincinnati Children’s Hospital Medical Center, maternal blood samples were obtained only at the time of delivery (n = 51). In contrast, at the University of Florida Health Women’s Health Center – Medical Plaza, maternal blood samples were collected both at a prenatal visit and again at the time of delivery (n =22).

### Extracellular Vesicle Isolation and Nanoscale Flow Cytometry (NanoFACS)

For extracellular vesicle (EV) isolation and cell-of-origin analysis, fresh whole blood was collected in EDTA tubes, centrifuged twice at 2500 × g for 15 minutes to obtain platelet-poor plasma (PPP), aliquoted (100 µL), and stored at −80°C. All samples were processed within 2 hours of collection and banked for batch testing. EVs were analyzed by nanoscale flow cytometry as previously described^6^. Antibodies (e.g., PLAP-APC, HLA-G-488; CD41-FITC, CD9-PE, CD63-PEcy7) were titrated (0.02–0.2 µg/mL) and incubated with platelet poor plasma. Samples were diluted 1:300 in 200 nm bead buffer (FACS-sorted polystyrene beads in 0.1 µm-filtered PBS), providing a relative size standard and plasma volume tested denominal reference and tested in triplicate experiments Acquisition was performed on a FACS Aria Fusion (BD Biosciences) with 70 µm nozzle and 0.1 µm-filtered sheath fluid at 70 psi. Instrument settings were optimized using Megamix beads (100–900 nm), and standard gates were applied across all batches. Data are reported as mean EVs per microliter of starting plasma using n=1000 200nm bead counts as volume control. EV identity and purity were validated using nanoparticle tracking analysis, cryo-electron microscopy (cryo-EM), and transmission EM (TEM) as previously described^6^.

### miRNA isolation and quantification

Total circulating and exosomal miRNAs were isolated from plasma samples using the Norgen Plasma/Plasma Circulating and Exosomal RNA Purification Kit (Slurry Format) (Norgen Biotek Corp.) following the manufacturer’s instructions. Briefly, 2 mL of plasma was combined with pre-warmed Lysis Buffer A and Slurry C2 supplemented with β-mercaptoethanol, mixed thoroughly, and incubated at 60 °C to ensure complete lysis. Ethanol was added to facilitate RNA precipitation, and the mixture was applied to Norgen mini spin columns for RNA binding. Bound RNA was subjected to on-column DNase I treatment using Norgen’s RNase-Free DNase I Kit (Cat# 25710) to eliminate genomic DNA contamination. Columns were then washed multiple times, and purified RNA was eluted in 50 µL of Elution Buffer. The resulting RNA samples were stored at –80 °C until use. RNA concentration and purity were assessed using a NanoDrop 1000 spectrophotometer (Thermo Fisher Scientific). To verify the presence and relative abundance of small RNAs, miRNA content was quantified using the Invitrogen Qubit microRNA Assay Kit (Cat# Q32280, Q32881) according to the manufacturer’s protocol. Synthesis of cDNA from purified miRNA was done using the TaqMan™ Advanced miRNA cDNA Synthesis Kit (Applied Biosystems), following the standard four-step enzymatic protocol provided by the manufacturer. Reverse transcription reactions were performed in a thermal cycler under standard cycling conditions for each enzymatic step (poly(A) tailing, adaptor ligation, reverse transcription, and miR-Amp preamplification). Quantification of individual miRNAs was carried out using TaqMan™ Advanced miRNA Assays (Applied Biosystems) specific for hsa-miR-29c-3p, hsa-miR-22b-3p, and hsa-miR-421. Expression levels were normalized to hsa-miR-19b-3p, which served as an endogenous control. Quantitative PCR (qPCR) was performed in triplicate for each target using QuantStudio3 (Applied Biosystems). Relative miRNA expression was calculated using the ΔΔCt method.

### Statistics

Statistical analyses were performed using two-way analysis of variance (ANOVA) and unpaired Student’s t-tests, as appropriate. Data are presented as mean ± standard deviation (SD). A p-value < 0.05 was considered statistically significant, unless otherwise stated.

## RESULTS

### Circulating total EVs are increased in maternal plasma of mothers carrying a fetus with a CHD

We investigated the impact of CHD on circulating EVs counts throughout gestation. The total extracellular vesicle count in plasma of control and CHD groups collected at the prenatal visit (16-24 weeks) and time of delivery (<37 weeks) obtained from NanoFACS. There were no significant differences in EVs abundance between control and CHD pregnancies at the prenatal visit. However, at delivery total EV nanocounts were significantly higher in the CHD cohort compared to controls (*p=0*.*0072*). Our results revealed that the increase in total EV abundance is associated with CHD (*p=0*.*0123*) rather than the time of collection, indicating a CHD associated alteration in circulating EV levels. Interestingly, there was no significant difference in levels of PLAP+, STB, or EVT derived EVs at the prenatal visit or at the time of delivery between control and CHD pregnancies (Figure 1B-D).

**Figure 1:**
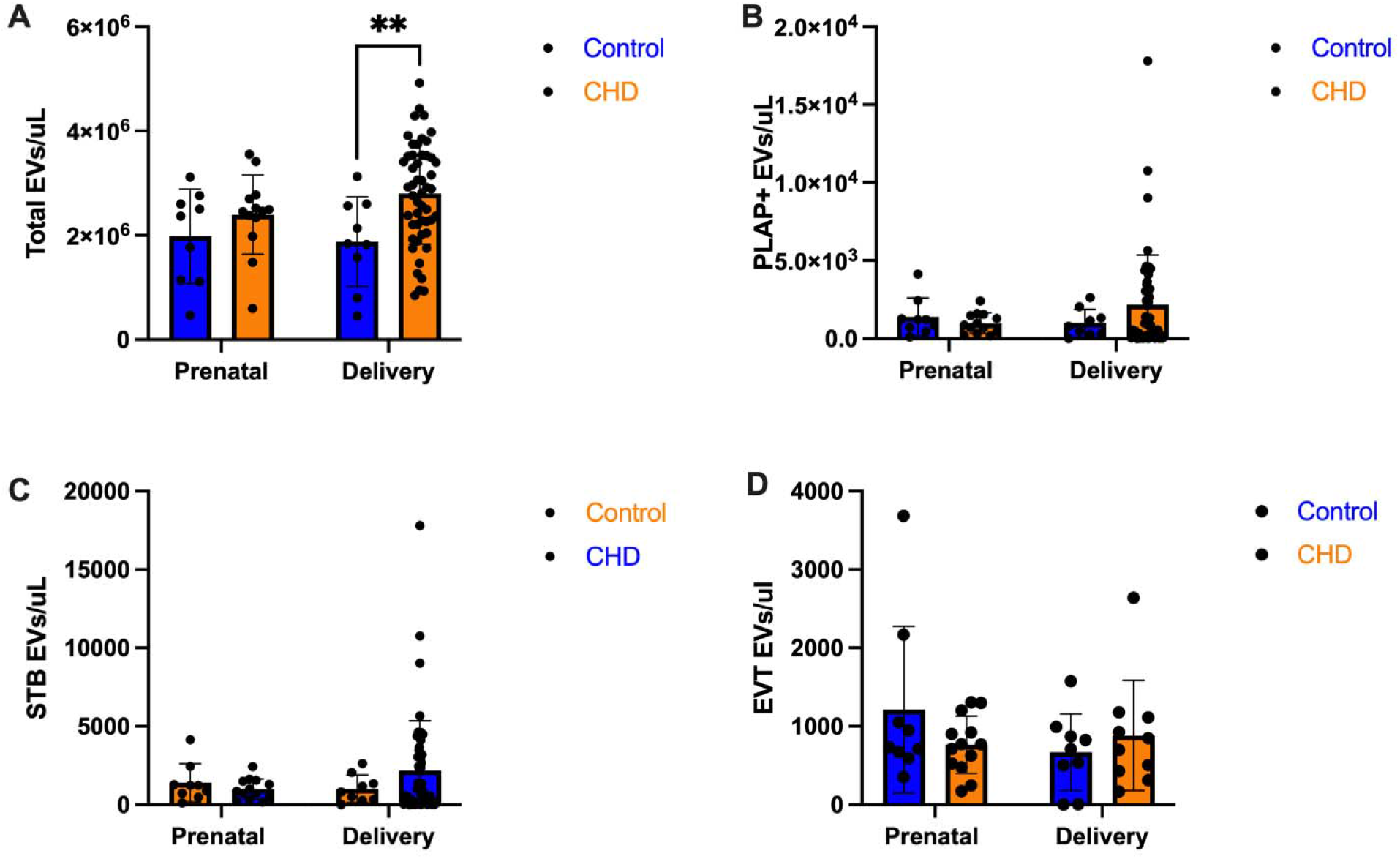
Quantification of EVs in maternal plasma from pregnancies with and without fetal CHD. **(A)** Total EV nanocount was significantly higher at the time of delivery in the CHD cohort (n=51) compared to control (n=9, p = 0.0072) pregnancies. **(B)** PLAP+ EV showed no significant difference between control and CHD pregnancies. **(C)** Syncytiotrophoblast (STB) derived EV showed no significant difference between control and CHD pregnancies. **(D)** Extravillous trophoblast (EVT) derived EV between control and CHD pregnancies. For each experiment a two-way ANOVA test was performed to determine significance. Values are represented as mean ± SD. **p<0.01. Prenatal refers to maternal plasma collected at 16-24 weeks gestation. EV, extracellular vesicle; CHD, congenital heart defects; PLAP, placental alkaline phosphatase; STB, syncytiotrophoblast; EVT, extravillous trophoblast.

### Activated platelet-derived EVs are increased at delivery in CHD pregnancies

We quantified activated platelet EVs in maternal plasma from control and CHD pregnancies at the prenatal visit and at the time of delivery (Figure 2). There was no significant difference in activated platelet-derived EVs between control and CHD pregnancies at the prenatal visit. However, at the time of delivery, the total nanocount of activated platelet-derived EVs wa significantly higher in the CHD cohort (p=0.010) compared to controls. Additionally, a significant increase in activated platelet-derived EVs was observed within the CHD cohort when comparing the prenatal visit to delivery (p=0.0051). However, we did not observe this same effect at the control cohort. We further revealed a significant interaction between CHD diagnosis and time of sample collection, indicating that the increase in activated platelet-derived EVs is dependent on both disease status and gestational age (p=0.0272).

**Figure 2:**
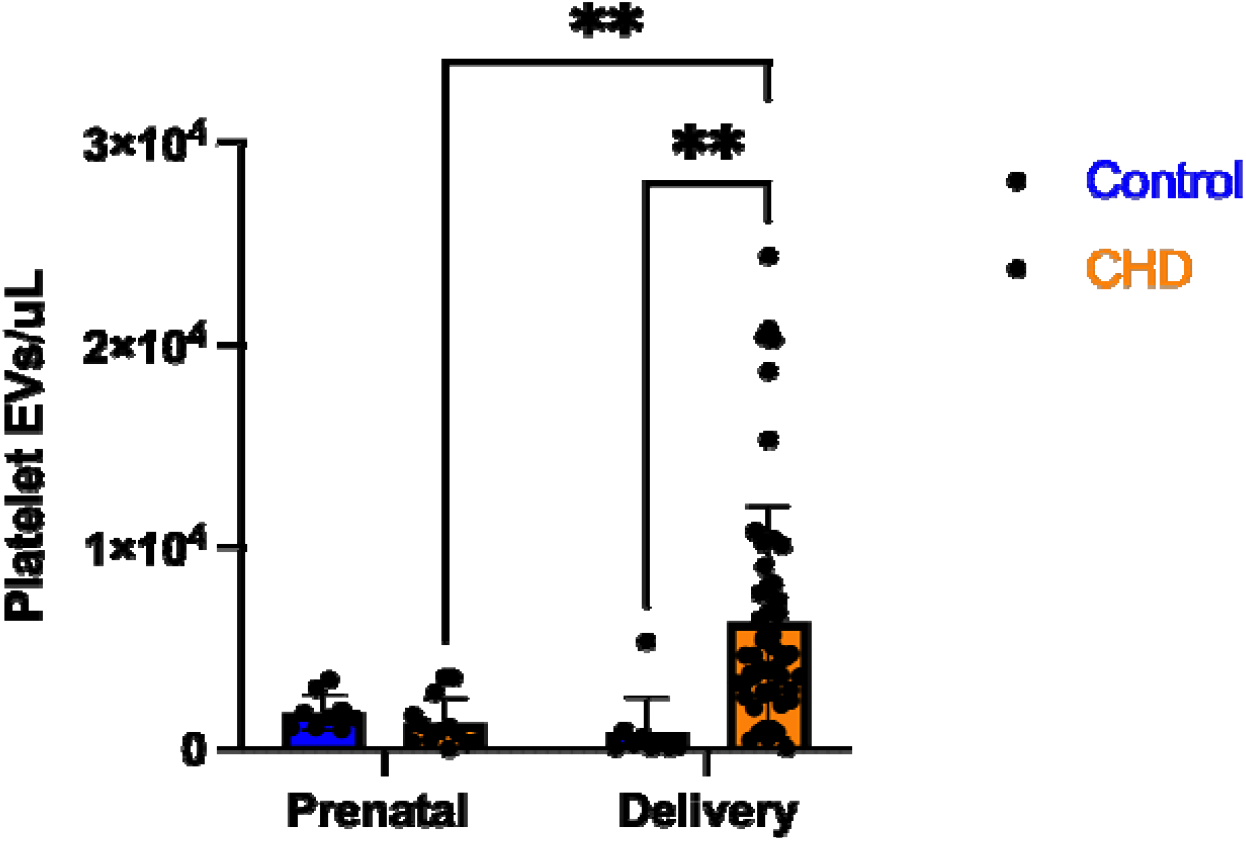
Quantification of activated platelet-derived EV in maternal plasma from pregnancies with and without CHD. **(A)** Activated platelet-derived EV nanocounts were quantified in maternal plasma collected at the prenatal visit and at the time of delivery from control and CHD pregnancies. At delivery, the total nanocount of activated platelet-derived EVs was significantly increased in the CHD cohort (n=51) compared to controls (n=9, p<0.01). For each experiment a two-way ANOVA test was performed to determine significance. Values are represented as mean ± SD. **p<0.01. Prenatal refers to maternal plasma collected at 16-24 weeks gestation. EV, extracellular vesicle; CHD, congenital heart defects.

### Activated platelet-derived EVs are elevated in cord blood at the time of delivery from CHD pregnancy

To assess whether CHD is associated with altered EV profiles in the umbilical circulation, we quantified total EVs and platelet-derived EVs in cord blood from control and CHD pregnancies. No significant differences in total circulating EV or endothelial EV counts were observed between control and CHD groups (Figure 3A-B). However, the total count of platelet-derived EVs was significantly higher in cord blood from the CHD cohort compared to controls (Figure 3B, p = 0.0389).

**Figure 3:**
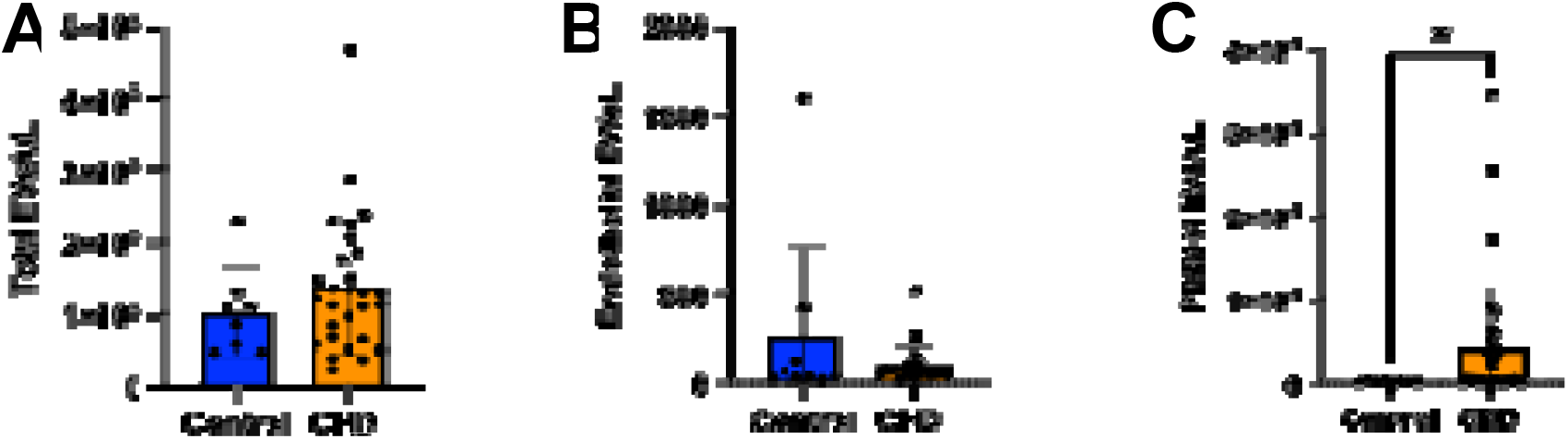
Quantification of EVs in cord blood from pregnancies with and without CHD. **(A)** Total EV nanocounts measured in cord blood did not differ significantly between control and CHD pregnancies. **(B)** Endothelial-derived EV nanocounts in cord blood plasma were not significantly different between groups. **(C)** Platelet-derived EV nanocounts in cord blood were significantly increased in the CHD cohort (n=28) compared to control (n=8). For each experiment a student’s t-test test was performed to determine significance. Values are represented as mean ± SD, *p<0.05. EV, extracellular vesicle; CHD, congenital heart defects.

**Figure 5.**
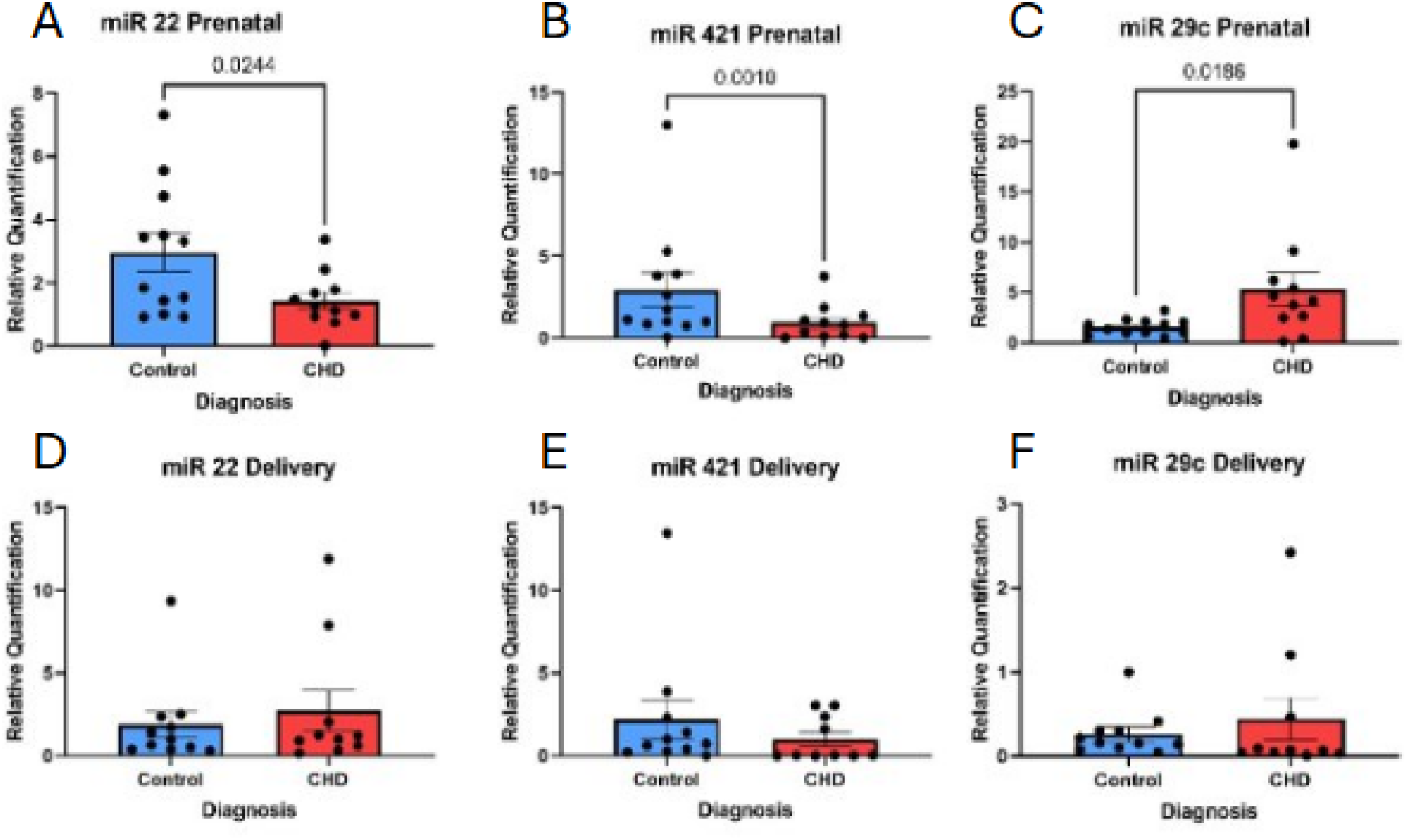
Differential miRNA expression in maternal plasma from pregnancies with and without CHD at prenatal and delivery. Relative expression levels of miR 22, miR 421, and miR 29c were quantified in maternal plasma from control and CHD pregnancies at the prenatal visit (A–C) and at the time of delivery (D–F). At the prenatal visit, miR 22 (A) and miR 421 (B) expression levels were significantly reduced in the CHD cohort compared to controls, while miR 29c (C) expression was significantly increased. No significant differences in the expression of miR 22 (D), miR 421 (E), or miR 29c (F) were observed between control and CHD pregnancies at the time of delivery. For each experiment a student’s t-test was performed to determine significance. Values are represented as mean ± SD. *P<0.05, **P<0.01. CHD, congenital heart defects; miR, microRNA.

### Differential expression of miR 22, miR 29c, miR 421 in plasma of mothers carrying a fetus with a CHD

To determine whether maternal circulating miRNA expression is altered in pregnancies affected by CHD, we quantified miR 22, miR 421, and miR 29c levels in maternal plasma at the prenatal visit and at the time of delivery using RT-qPCR. At the prenatal visit, maternal plasma levels of miR 22 and miR 421 were significantly reduced in CHD pregnancies compared with controls (miR 22: p = 0.0244; miR 421: p = 0.0010; control, n = 12; CHD, n = 11). In contrast, miR 29c expression was significantly increased in maternal plasma from the CHD cohort relative to controls at the prenatal visit (p = 0.0186; control, n = 12; CHD, n = 11). At the time of delivery, no significant differences were observed in the expression levels of miR 22, miR 421, or miR 29c between CHD and control pregnancies.

## DISCUSSION

This is the first study to identify the cellular sources of circulating EVs in pregnancies complicated by CHD. The focus of this study was to quantify circulating EVs and miRNA during 2^nd^ trimester and at term in the plasma of mothers carrying fetuses with CHD and in umbilical cord plasma at delivery. It is known that as pregnancy progresses, the secretion of EVs increases^13^. Furthermore, in the third trimester of pregnancy, the concentration of EVs increases by 3-fold, largely due to the production of platelet and endothelial derived EVs^14^. Our results demonstrated a significant increase in total circulating EVs in the CHD cohort at the time of delivery compared to control pregnancies. This elevation in total EV is likely due to the overall increase in platelet derived EVs. Furthermore, we did not observe a difference in the PLAP+, STB or EVT derived EVs in maternal plasma at either point in gestation in the CHD cohort.

Platelets contribute to hemostasis by supporting blood clotting and play a crucial role in modulating the immune environment and promoting tissue repair^15^. Platelet-derived EVs are released upon platelet activation including vascular injury or inflammation. We observed that EVs derived from activated maternal platelets were significantly upregulated at the time of delivery in the CHD cohort. Recent evidence has demonstrated that CHD pregnancies have an increased incidence of maternal malperfusion^16^ which is a consequence of defective spiral artery remodeling by EVT, suggesting that as pregnancy progress, the expansion of maternal blood volume ^1^ near term may cause shear stress on the maternal vasculature and result in vascular dysfunction and inflammation. This dysfunction may trigger platelet activation, leading to the systemic release of pdEVs potentially containing pro-inflammatory^19^ and pro-thrombotic cargo^20,21^. The rise in pdEVs observed at delivery is similar to that seen in preeclamptic patients^22^. Therefore, these findings suggest that the increase in pdEVs near delivery may represent a maternal adaptive response to vascular stress or injury, which is seen in pregnancies affected by both fetal CHD and preeclampsia. This response could be a protective mechanism to promote hemostasis and prevent blood loss during childbirth^23^, particularly since mothers who carry a fetus with CHD have an elevated risk of hemorrhage at delivery^24^.

In this study, we analyzed total circulating miRNAs with the understanding that the majority of circulating miRNAs are found within EVs that could be released from any placental or maternal cell type. Consistent with previous studies that reported higher expression of miR-29c in the plasma of mothers carrying fetuses with a VSD diagnosis^12^, we observed higher levels of miR-29c in our varied cohort of CHD pregnancies at the prenatal visit compared to controls. miR-29c regulates transcripts involved in apoptosis, extracellular matrix (ECM) remodeling, and cell adhesion. Studies have shown that miR-29c is a negative regulator of matrix metallopeptidase-2 and integrin β1^25^, both essential for ECM remodeling and effective EVT invasion. Additionally, studies show that overexpression of miR-29c promotes apoptosis ^26,27^, suggesting that increased miR-29c in CHD pregnancies may be a sign of stress and potentially apoptotic signaling. Our findings also suggest the production of this miRNA by EVTs results in impaired maternal spiral artery remodeling and potentially contributes to placental insufficiency during the prenatal visit in CHD pregnancies.

Our results show that miR-22 is significantly decreased at the prenatal visit but turn to baseline at the time of delivery. There is evidence to suggest that, in inflammatory states, the NF-kB pathway is activated in endothelial cells, suppressing miR-22 expression^28^. This reduction in miR-22 leads to activation of the AKT/mTOR signaling pathway, which is essential for normal endothelial function^29^. Studies using hypertensive rat models show that these effects can be reversed in maternal exercise by inhibiting the growth factor signaling pathway to improve vascular remodeling^30^. In healthy endothelium, AKT phosphorylates endothelial Nitric Oxide Synthase (eNOS) to increase Nitric Oxide (NO) production. NO is essential in relaxing smooth muscle and initiating vasodilation in maternal arteries, reducing vascular resistance and allowing adequate blood flow into the placenta^31,32^. miR-22 plays roles in glucose metabolism and cellular stress responses by negatively regulating the GLUT4/PI3K/AKT pathway in placental tissue in patients with gestational diabetes^33^. Additionally, studies have shown that higher blood glucose levels in pregnancies correlate with a higher risk for fetal CHD, even in mothers who are not diabetic^34^. A reduction in miR-22 may indicate a maternal inflammatory response that regulates glucose signaling in the placenta, potentially contributing to an increase in placental glucose uptake. In our cohort, miR-421 expression was significantly decreased at the prenatal visit, followed by a return to baseline levels by term. Previous studies report overexpression of miR-421 in right ventricular tissue of infants with TOF, where it is associated with altered cardiac remodeling^35^. Previous studies have demonstrated that miR-421 expression is decreased in STB-EV maternal serum from preeclamptic patients compared to controls^36^. Although our study did not observe significant differences in total STB-EV abundance between CHD and control pregnancies, the reduction in circulating miR-421 suggests that EV cargo composition may still be altered in CHD pregnancies. Predicted and CLIP-seq-supported targets of miR-421 include genes involved in oxidative stress responses, cellular metabolism, cytoskeletal remodeling, and energy sensing, including PRKAA2 and AK4^36^. PRKAA2 encodes a catalytic subunit of AMPK, a central regulator of cellular energy homeostasis and oxidative stress adaptation^37,38^, while AK4 is associated with mitochondrial stress responses and metabolic regulation^39,40^. Together, these findings suggest that reduced miR-421 expression may reflect altered placental EV cargo signaling in CHD pregnancies, potentially contributing to placental impaired metabolic adaptation in trophoblast.

In conclusion, this study is the first to characterize the cellular sources of EV and circulating miRNA levels in pregnancies complicated by CHD. We demonstrated that CHD pregnancies are associated with an altered EV and miRNA profile in a gestational age-dependent manner. We show that there is a difference in total EV and platelet derived EVs but not PLAP+ derived EVs. Nevertheless, these findings support the concept that CHD pregnancies are characterized by gestational age-specific changes in EV release and miRNA signaling. These changes may reflect altered placental and maternal vascular remodeling or a metabolic adaptation in CHD

## Notes

### Competing Interest Statement

The authors have declared no competing interest.

